# Backward evolution from gene network dynamics

**DOI:** 10.1101/369371

**Authors:** Merzu Kebede Belete, Daniel A. Charlebois, Gábor Balázsi

**Affiliations:** The Louis and Beatrice Laufer Center for Physical and Quantitative Biology, Stony Brook University, Stony Brook, New York, 11794-5252, USA; Center for Systems and Computational Biology, Caner Institute of New Jersey, Rutgers University, New Brunswick, New Jersey, 08901, USA; Department of Biomedical Engineering, Stony Brook University, Stony Brook, New York, 11794-5281, USA

**Keywords:** Evolution, gene regulatory network, Markov process, Monte Carlo simulation

## Abstract

Gene expression is controlled by regulator genes that together with effector genes form gene regulatory networks. How mutation in the genes comprising gene regulatory networks influences cell population dynamics has not been adequately investigated. In this study, we develop mathematical models to study how a mutation in a regulator gene that reaches the effector gene with a time delay affects short-term and long-term population growth. Using theory and experiment, we find a paradoxical outcome of evolution where a mutation in a regulator gene leads to an interaction between gene regulatory network and population dynamics, causing in certain cases a permanent decrease in population fitness in a constant environment.

**Significance Statement:** The properties of a cell are largely the products of its proteins, synthesized at rates depending on the regulation of protein coding genes. Single-cell measurements show that genetically identical cells can differ radically in their protein levels, partially due to the random production and degradation of proteins. It is currently unknown how mutants arise and spread in populations affected by such biological variability. We use computer simulations and evolution experiments to study how a mutant spreads in a population that carries a synthetic drug resistance gene network. Our results show for the first time a paradoxical outcome of evolution, where an initially beneficial mutation can interact with gene regulatory network dynamics and cause a permanent decrease in population fitness in the same environment.

## 1 Introduction

An organism’s genome encodes a large number of intertwined genes that form gene regulatory networks (GRNs). The products of these genes enable a number of cellular processes like stem cell differentiation (Balázsi et al, 2011; Janga & Collado-Vides, 2007; Nevozhay et al, 2012), drug resistance (Balaban et al, 2004; Charlebois et al, 2011a; Nevozhay et al, 2012), and metabolic activity (Dykhuizen et al, 1987; Glanemann et al, 2003; O’Brien et al, 2013). The product of a regulator gene (a gene whose expression product regulates the transcription of at least one gene in the network) can control the activity of a number of downstream regulator genes, effector genes (outputs of a GRN that influence cellular fitness), or a combination of both genes (Alon, 2007; Shen-Orr et al, 2002). The level and temporal pattern of effector gene products in the cell are important not only for cellular fitness (Levy et al, 2012; Nevozhay et al, 2012), but also influence population dynamics (Fraser & Kaern, 2009; Nevozhay et al, 2012).

Despite the fundamental importance of GRN evolution in biology, we have only begun to understand how they are rewired to adapt to environmental challenges (Bornholdt, 2001; Cohen et al, 2008; Dekel & Alon, 2005; González et al, 2015; Rancati et al, 2008) and how rewiring affects GRN and fitness dynamics. GRNs are not static entities, but rather they can evolve by adding or removing genes, or by altering the strength of connections between genes (Crombach & Hogeweg, 2008). Selective pressures act on gene networks to optimize their activity levels and confer a fitness advantage to the host organism (Kellis et al, 2003). For example, GRN evolution may occur through tuning the strength of negative-feedback regulation (Peng et al, 2015; Prud’homme et al, 2007), or by mutation both within and outside of a stress-response network (González et al, 2015).

Stressful environments can change the wiring of a gene regulatory network by selecting for mutations that lead to different dynamics in gene expression (González et al, 2015). A quantitative understanding of the effect of mutation on naturally occurring gene networks is difficult due to the high degree of interconnectivity and dependence between genes or networks in a genome (Maynard et al, 2010). As a result, little is known about how mutations in gene networks percolate and affect long-term population fitness. Synthetic gene circuits (Elowitz et al, 2002; Hasty et al, 2001; Nevozhay et al, 2013) are ideal to study evolution in a controlled manner, as they typically do not interact with the host genome, consist of well-characterized components, and greatly reduce the parameter space of evolution (González et al, 2015).

Mutations in a regulator gene, many of which are likely to be deleterious by causing disruption of existing regulatory interactions, can influence the performance of a gene network. For instance, mutations can affect regulatory interactions by creating a new mutant protein which (i) may recognize a different DNA sequence (ii) may not bind to the DNA site of the effector, (iii) may not associate with the inducer if the regulation process is inducible, or (iv) binds to the inducer but is unable to activate the gene cascade. The effects of mutation take time to appear at the phenotype level, as the effect of mutation must percolate through regulatory layers in the network (including mRNA and protein synthesis, as well as additional regulators if the effect of the activator gene is indirect). The time scales of these intracellular processes may be comparable to the time scale of dynamics at the population level, which may have unexpected consequences for evolution.

We develop mathematical models and perform stochastic simulations to study the interplay between mutation in a gene regulatory network cascade and population dynamics. More specifically, we investigate how the time-delayed effect of mutation in a regulator gene on an effector gene predicts the population fitness. We mathematically explore the parameter space of the mutant’s cellular fitness and the time delay to determine long-term population fitness. We find that when the time delay of mutation to reach in the effector gene is small, the mutant subpopulation always collapses. For a long time delay and sufficiently large initial cellular-fitness benefit from a reduction in rtTA toxicity or “squelching” (Baron et al, 1997), the mutants sweep the population which ultimately results in lower population fitness compared to the wild-type (ancestral) cell population. This is due to an increase in antibiotic susceptibility, as antibiotic resistance proteins (which are no longer being produced in mutant cells) decay. In an intermediate relationship of delay and cellular-fitness benefit the mutant subpopulation undergoes one of two stochastic fates: either collapse or sweep of the population. Finally, an evolution experiment was conducted on budding yeast *Saccharomyces cerevisiae* carrying a well characterized synthetic stress response gene circuit (Nevozhay et al, 2012) which supports our model predictions.

## 2 Results and Discussion

### 2.1 Mutation and fitness dynamics in genetic cascades

To generally study the effect of mutation of a regulator gene in a GRN on cellular fitness, we first consider a model of an *n* gene cascade where *n* − *1* are the regulator genes and the other gene is an effector gene [Figure 1(A)]. The protein of gene *i* positively regulates its adjacent gene *i* + *1*. We assumed that the regulatory proteins are cellular fitness inhibitors due to the toxicity costs associated with induction (Baron et al, 1997), while the effector protein enhances cellular fitness in the presence of a stressor (Nevozhay et al, 2012). When a mutation occurs in a regulator gene, it affects that regulator’s DNA sequence. The new DNA sequence is a template for producing mutant mRNA, which is translated into a mutant regulator protein which is not toxic to the cell and incapable of activating gene expression [Fig. 1(B)]. An intact protein pool and non-mutant mRNA can temporarily mask the mutation. The effect of mutation will gradually appear as a change in cellular fitness associated with lower levels of intact regulator protein, and will percolate through the layers of the network, until it finally reaches (reduces induction) the effector gene and the cells succumb to the effects of the drug.

**Figure 1:**
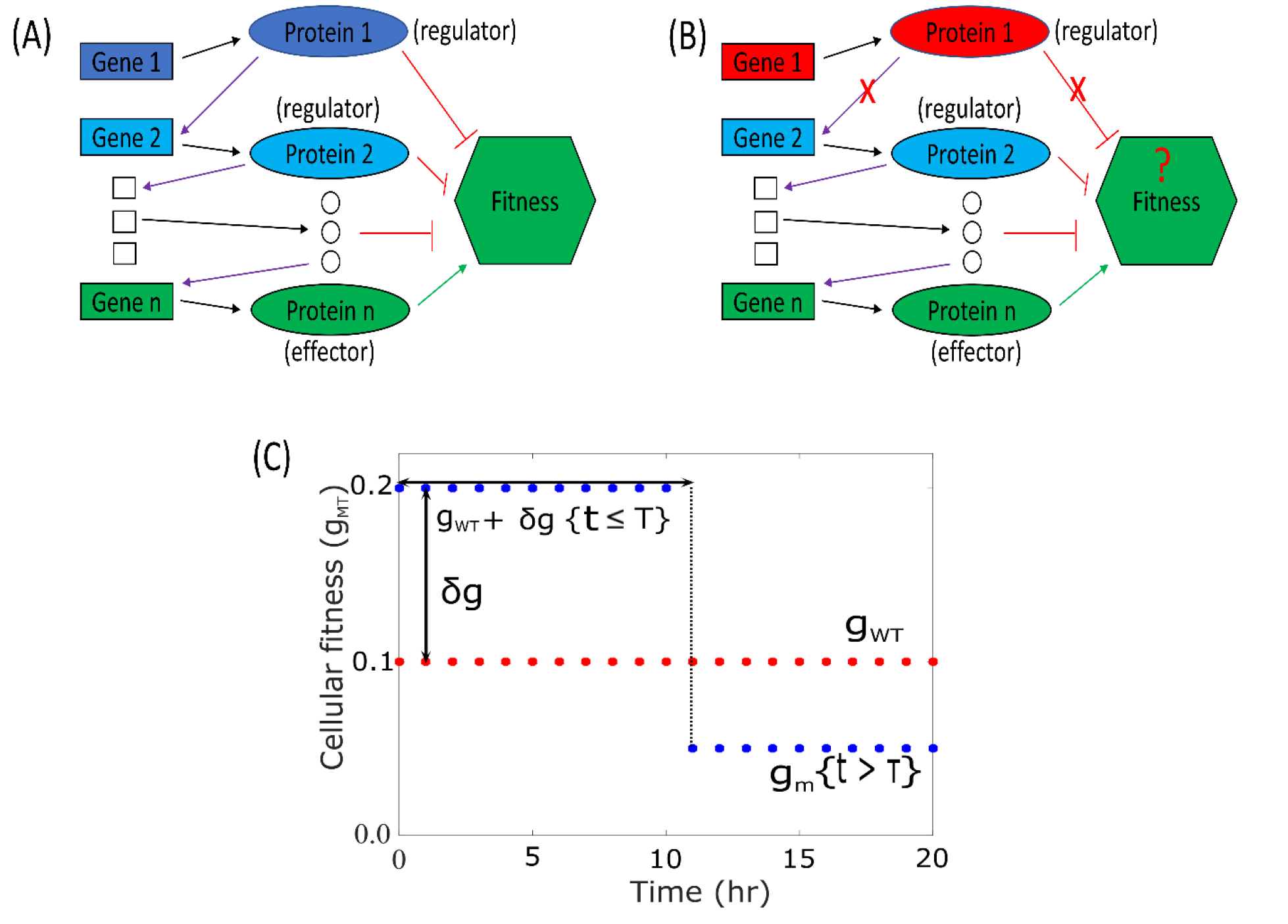
Schematic of an n-gene network cascade for wild-type and mutant cells. **(A)** A gene cascade in a wild-type cell: gene *i* produces a protein *i* (regulator) which activates gene *i* + *1*. Here all the regulators are assumed to inhibit fitness. The protein *n* (effector) enhances cellular fitness. **(B)** A gene cascade in a mutant cell: here a mutation in gene 1 produces a mutant regulator protein which does not activate the circuit. The effect of the time delay (τ) and mutation in gene 1 is represented by ? and X, respectively. **(C)** The time series of the cellular fitness of the mutant (*g_MT_*) with a τ and fitness benefit (*δ_g_*). The final mutant cell fitness (*g_m_*) in this case decreases to a level below wild-type cell fitness (*g_WT_*) after the time delay. In (A) and (B), red rectangles and ellipses respectively represent mutated genes and proteins, black arrows denote transcription/translation, purple arrows denote transcription factor activation, and green arrows and red blunt arrow denote positive and negative effects on fitness, respectively.

To investigate the dynamics of a single mutation on population fitness, we assume the total mutant single-cell fitness in a stressful environment is due to the effect of both regulator and effector protein levels (Nevozhay et al, 2012; Yeh et al, 2009). After a mutation occurs in a regulator gene, the mutant cellular fitness level increases by *δ_g_* (cellular fitness benefit) for a length of time (described by a time delay, τ), due to a decrease in intact regulator protein level and its associated cytotoxicity [Fig. 1(C)]. This is followed by a reduction in fitness due to a decrease in the level of the drug resistant effector protein. The cellular fitness of the mutant, *g_MT_(t)*, can be approximated by first a stepwise increase, and then by a stepwise decrease,

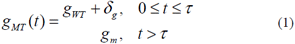

where *g_MT_(t)* and *g_WT_* are the cellular fitness of the mutant and wild-type cell, respectively, and *g_m_* is the cellular fitness of the mutant after the time delay.

### 2.2 Stochastic fate of mutants

We ran stochastic simulations of a dividing cell population while keeping the number of cells constant. The simulations were initiated with a single mutant in a wild-type cell population, and the dynamics of both the mutant and wild-type cells were investigated. The results in Figure 2 show the time-dependent extinction and fixation histograms, fraction of mutant cells (*f*), and time course of the population fitness. In a growing mixed cell population, beneficial mutations can be rapidly lost due to genetic drift during the process of random sampling (Kimura, 1962). If the mutant escapes drift, the mutant population increases its frequency and undergoes three possible fates as we change either *δ_g_* and keep τ constant, keep *δ_g_* constant and vary *τ*, or vary both parameters. The histograms in Figure 2(A, D, & G) show the steady state probability distributions for the time of extinction and fixation of mutant for *δ_g_* = [0.04, 0.075, 0.11], respectively. The mutant can vanish due to genetic drift at an early time, shown as a Poisson-like distribution in Figure 2(A, D, & G), or go extinct due to fitness loss after τ [Fig. 2(A & D)]. The mutant subpopulation can also take over the population, if escapes genetic drift and has a sufficient fitness benefit in the early time of *t* ≤ τ [Fig. 2(D & G)].

**Figure 2:**
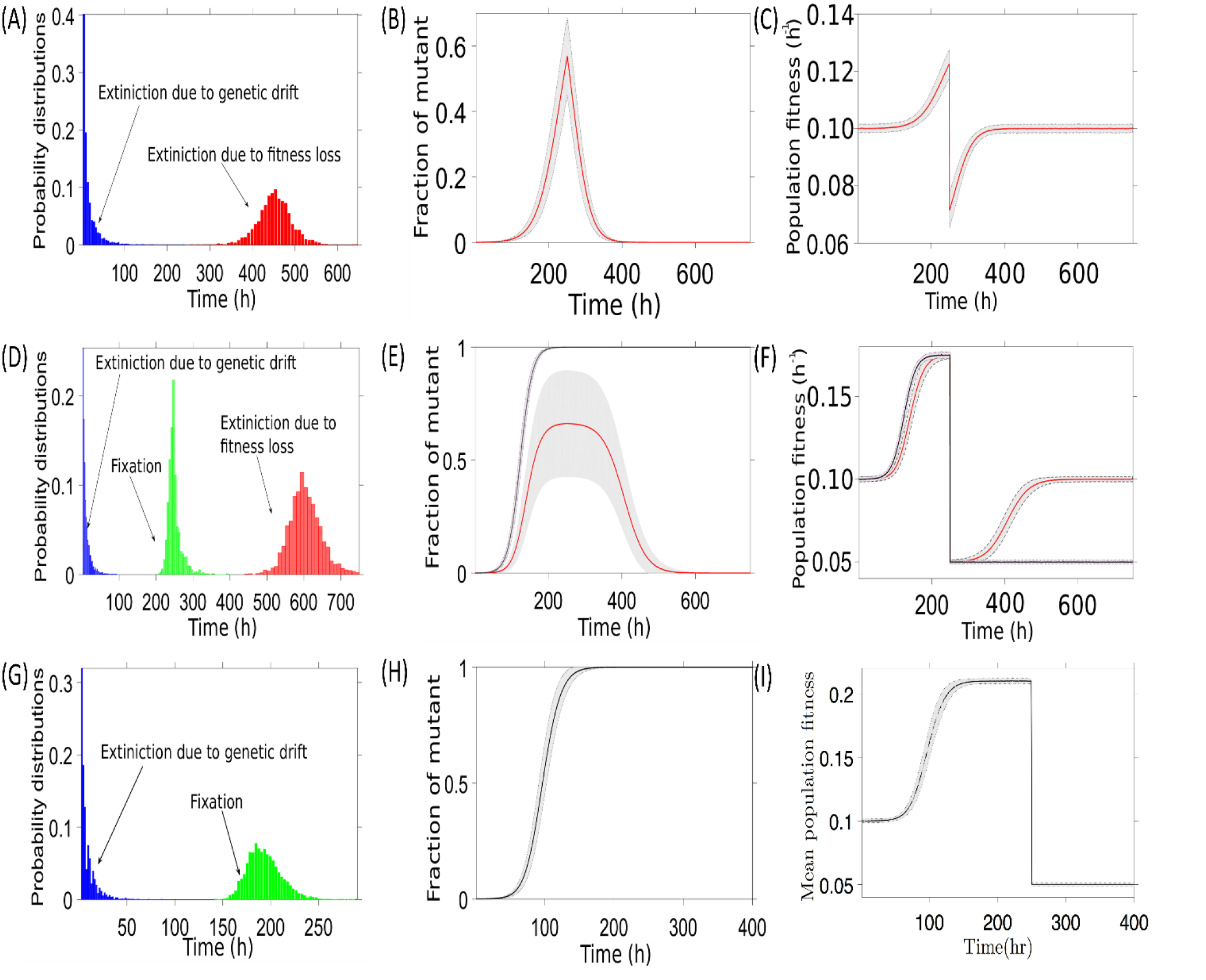
Time courses of mixed population fitness, fraction of mutants, and fixation-extinction histograms. **(A-C)** The dynamics of mutants with a low fitness benefit δ*_g_*= 0.04 h^−1^. The mixed population fitness initially increases with the increasing fraction of mutants which confers a fitness advantage (C), then eventually the mutant population collapses due to a loss in fitness advantage (B). **(D-F)** In the intermediate fitness benefit *δ_g_* = 0.075 h^-1^ parameter regime, the mutant subpopulation undergoes one of two stochastic fates: either it sweeps the population (*f* = 1) or it collapses (*f* = 0) (E) with lower or wild-type long-term population fitness, respectively (F). **(G-I)** For a high fitness benefit *δ_g_* = 0.11 h^−1^, the mutation which escape extinction due to genetic drift sweeps the population (H) with a lower permanent population fitness (I). (A, D, and G) Histograms that show the time of fixation and extinction probability distributions of mutants. The mutants can vanish due to genetic drift at the early time with a Poisson-like distribution (shown in blue) or lost after delay τ due to the deleterious properties of the mutant (shown in red). The mutants also have a chance take over the population (shown in green) due to escape in genetic drift and fitness benefit when *t* ≤ τ. Here, parameters were set to τ = 250 h, *g_WT_* = 0.10 h^−1^ and *g_m_* = 0.05 h^−1^ and were obtained from the phase space shown in Figure 3(A). Shaded areas denote standard deviation.

For a low-cellular fitness benefit, the mutant subpopulation always collapses (*f* = 0) [Fig. 2(B)] and the long-term population fitness returns to that of the wild-type population [Fig. 2(C)]. When the cellular fitness benefit is high, the mutant cells always sweep the population (*f* = 1) [Fig. 2(H)] and long-term population fitness decreases below that of the wild-type population [Fig. 2(I)]. For an intermediate cellular fitness benefit, the mutant cells have two stochastic fates (*0 < f < 1*) [Fig. 2(E)]: they either take over the population, lowering population fitness, or the mutant subpopulation collapses and population fitness returns to its initial value (wild-type growth rate) [Fig. 2(F)].

Next, we investigate the fate of mutants carrying the gene network architecture shown in Figure 1(B). The increase in the number of mutant cells in nutrient rich environment is modeled as an exponential process. The fraction of mutant cells, *f*, at *t* = τ starting from a single-mutant cell from a population of *N_0_* wild-type cells at *t* = 0 is described by,

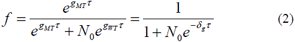

Rewriting the above equation yields,

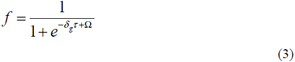

where Ω=*ln(N_0_)*. The above equation allows us to relate *δ_g_* and τ to the fraction of mutant cells in the population. Equation 3 is similar to the logistic equation (Verhulst, 1845; Verhulst, 1847), as *f* levels off after an exponential growth period as δ_*g*_τ increases.

To test the robustness of the discrete-state model, we ran simulations varying both the time delay and the cellular fitness benefit (Fig. 3). The mutant population dynamics undergoes a phase transition as we vary τ and δ*_g_*[Fig. 3(A & B)]. For a short time delay or low cellular fitness benefit, the mutant population always collapses (*f* = 0). For a long time delay or high cellular fitness benefit, the mutant cells always sweep the population (*f* = 1). As before, there is regime of phase space where fate of mutant cells is stochastic (*0 < f < 1*).

**Figure 3:**
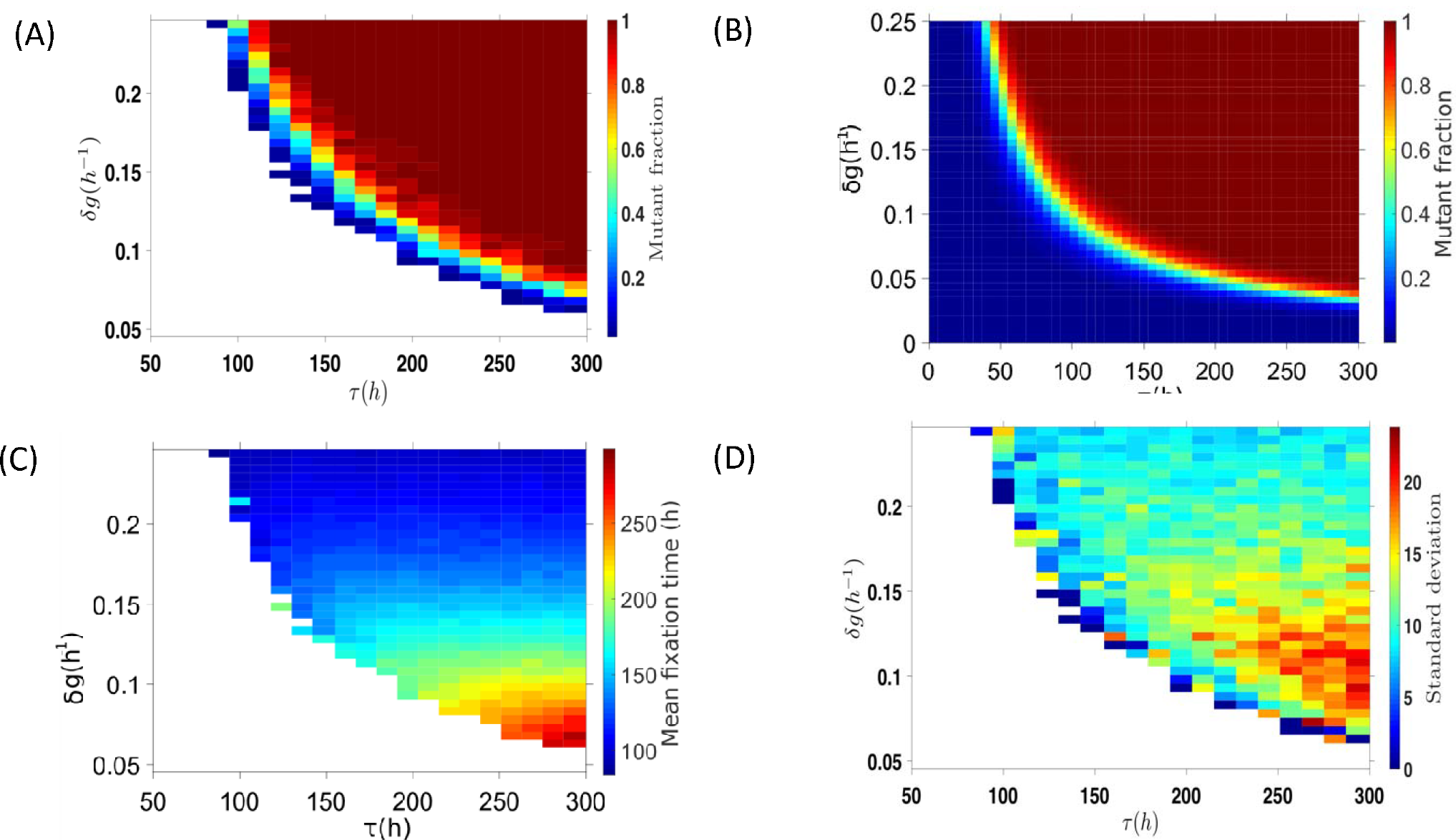
Effect of mutation benefit and time delay on mutant fraction and fixation time in a mixed cell population. **(A)** Heat map of the mutant fraction that sweeps the population as a function of cellular fitness benefit (*δ_g_*) and time delay (τ). The color bar shows the fraction of mutants that sweep the population (number of mutants which fix in the population divided by number of mutants which fix plus the number of mutants which go extinct due to fitness loss). **(B)** Analytic solution corresponding to (A) obtained from Equation 2. **(C)** Mean-fixation time of the mutants. **(D)** Standard deviation of the fixation time of the mutants. White regime (not shown on color bars) indicates parameter regimes where the mutant subpopulation went extinct.

A mutant which appears in a simulated fixed-size population will either fix in the population by escaping genetic drift and outcompeting wild-type cells, or it will go extinct. Figure 3(C & D) shows the mean and standard deviation of the fixation time of mutant which escapes genetic drift, respectively. As *δ_g_* gets smaller, both the mean and the standard deviation of the fixation time increase.

### 2.3 Mutation and fitness dynamics in PF yeast cell population

So far we assumed that there is no phenotypic variation between cells in the population. However, genetically identical cells can differ significantly in their gene-expression level (Acar et al, 2008; Charlebois & Kaern, 2012; Elowitz et al, 2002), which generates cellular fitness diversity or “fitness noise” (Levy et al, 2012; Nevozhay et al, 2012) within the population. To address such variations, we developed a coarse-grained stochastic model by dividing up the population gene expression profile into discrete cellular states (phenotypes). In each state, cells have specific cell division rate and can randomly switch into neighboring states due to a change in gene expression. We assume that the cells are randomly switching back and forth between adjacent states, giving rise to a purely Markovian process (van Kampen, 1992). The gene expression profile of a synthetic inducible positive feedback (PF) gene network [Fig. 4(A)], a common motif in natural transcriptional networks (Mitrophanov & Groisman, 2008), can be discretized into low-expressor (state 0), intermediate-expressor (state 1), and high-expressor (state 2) states where cells in the population can randomly switch states (see SI Section 3.2). We first calibrated this model using the location of the fitness peak in fixed drug and varying inducer conditions (González et al, 2015) and reproduced experimental results (Nevozhay et al, 2012) in a growing cell population that carries this gene circuit. We study the effect of time delay needed for a mutation in the regulator gene (rtTA) [Fig. 4(B)] to reach the resistance-reporter gene (yEGFP::zeoR), and affect the dynamics of the phenotypic switching population using both the coarse-grained model and a fine-grain population dynamics algorithm (PDA) (Charlebois et al, 2011b) (see SI Section 3.3).

**Figure 4:**
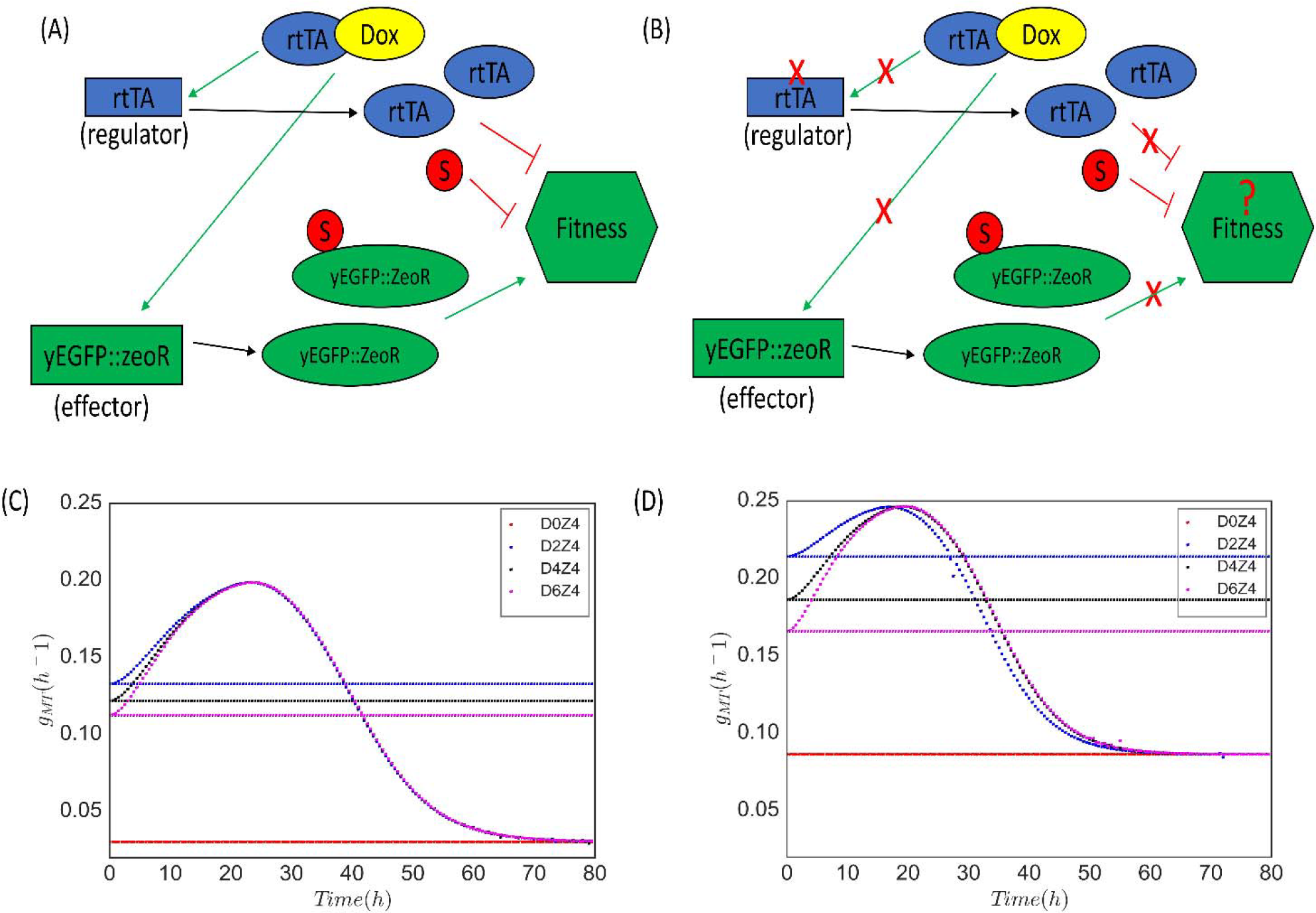
Diagram of the positive feedback gene construct for wild-type and mutant cells. The construct consists of a regulator gene (*rtTA*) and an effector gene (*yEGFP::zeoR*). Doxycycline (Dox) is the inducer and the drug Zeocin is the stressor (S). **(A)** For wild-type cells, Doxycycline enters through the cell membrane and binds to the regulator forming a complex *dox−rtTA*; the complex positively activates both the regulator and the effector genes. The effector protein protects yeast cells from the stressor (Zeocin) by forming a protein-stressor complex. **(B)** For mutant cells, the new regulator protein does not activate the circuit in four possible scenarios (see main text). However, there are sufficiently large numbers of intact protein which keep the circuit activated for a time duration *τ*. The Xs denote a missing interactions due to mutation of the *rtTA* gene and ? the unknown effect of the time delay on fitness. The green and black arrows represent activation and production, respectively, and the red flat head arrows represent fitness inhibition. **(C)** The cellular fitness of the mutant for different concentration of Doxycycline and a fixed Zeocin concentration (2 mg/ml). Mutant cellular-fitness starts from the wild-type (solid horizontal colored line for each condition) and increases and then decreases due to a decrease in effector protein concentration. The final cellular fitness of the mutant is lower than the wild-type cellular fitness. The time delay *τ* is defined as the time required for the mutant cellular fitness crosses the corresponding wild-type cellular fitness.

The regulator’s gene expression in the PF gene circuit is activated by adding an inducer (Doxycycline) (Nevozhay et al, 2012). The regulator gene controls the expression of an effector protein which provides resistance to the drug Zeocin. The effector protein consists of the fluorescent reporter yEGFP fused to the drug resistance gene zeoR (*yEGFP::zeoR*). Activating the circuit is both toxic and beneficial to the cell in the drug environment, as expressing the regulatory gene causes toxicity by squelching but protects the cell from the drug by activating *yEGFP::zeoR*. We note that cellular memory of the high expression state in the PF circuit was estimated to be in excess of 283 hours (Nevozhay et al, 2012).

In the presence of an inducer, the cell can mutate to overcome the effect of the inducer and consequently increase cellular fitness (González et al, 2015). However, after a time delay due to the decrease in regulator protein level, the level of drug resistance effector protein also decreases. Therefore, in the presence of both stressor and inducer, the mutant cell has a temporary fitness advantage for the time duration τ while it degrades the regulator protein. When the mutant cells start running out of the effector protein, we expect the mutant cellular fitness to drop [similar to Figure 1(C)]. The cellular fitness for each state was calculated from simple biochemical considerations of the PF gene circuit for both wild-type and mutant cells (see SI Section 4). The switching rates for the three-state PF cells were inferred from an earlier two-state model (Nevozhay et al, 2012), by matching the known gene distribution in the experiment. For the mutant cells, the switching of states is always in one direction from state 2 to state 1, and eventually to state 0 (absorbing state), with the switching rate equal to the cellular fitness of the mutant.

We then used our model to investigate the effect of increasing Doxycycline on the time delay for fixed Zeocin concentrations. Here DiZj denotes experimental conditions, where Di and Zj indicate the concentrations of the inducer Doxycycline (μg/ml) and the drug Zeocin (mg/ml), respectively. We found that when Zeocin was fixed at 4 mg/ml, τ increased with Doxycycline concentration [Fig. 4(C)]. A population crash can only occur if the fitness of the mutant falls below that of the wild-type, and when *δ_g_* is sufficient large or τ sufficiently long. The model therefore predicts that a fitness crash is most likely to occur in the D4Z4 or D6Z4 conditions. Similar results were found when Zeocin was fixed to 2 mg/ml, though with shorter τ s due to the increased effects of the drug [Fig. 4(D)].

Similar evolutionary dynamics were found for the PF gene circuit with the previous an *n*-gene network cascade and a three-gene cascade using the fine-grain population dynamics algorithm. Namely, the time delay of mutation to percolate through the PF gene circuit can affect the outcome of the PF cell population (see SI Section 3.3). The fine-grain PDA accounts for the processes of probabilistic cell division, stochastic switching of phenotypes, and uses a constant-number Monte Carlo technique to accurately simulate the statistics of an exponentially growing cell population (Charlebois et al, 2011b; Khalili et al, 2010; Mantzaris, 2006; Mantzaris, 2007). Evolutionary dynamics are generally assumed to push the population towards a peak in the fitness landscape [for a review see (de Visser & Krug, 2014)]. However, here we find that the mutations in PF cells interact with GRN dynamics and give rise to a permanent decrease in population fitness in the same environment, thus representing move towards a valley in the fitness landscape.

### 2.4 Evolution Experiments

We experimentally tested the predictions of our model using *Saccharomyces cerevisiae* carrying a synthetic two gene positive feedback (PF) gene circuit (Fig. 4) (see SI for experimental procedures). This gene network architecture is found in natural stress resistance networks, such as the yeast pleiotropic drug resistance network (Charlebois et al, 2014; Delahodde et al, 1995), the multiple antibiotic resistance network in Escherichia coli (Garcia-Bernardo & Dunlop, 2013) (nested with negative feedback) or implicitly, as growth mediated feedback (Deris et al, 2013), and was previously implemented in a synthetic gene circuit (Nevozhay et al, 2012). The synthetic PF gene circuit allows for separate control of the stress and the response by adjusting inducer and drug concentrations, respectively. Importantly, it is known that the PF circuit can evolve by intra- and extra-circuit mutations that modulate gene expression depending on the stress-response balance (González et al, 2015).

Zeocin is a broad-spectrum antibiotic that causes cell death by passively diffusing through the cellular membrane and inducing double-strand DNA breaks. When Zeocin is unbound with the resistance protein, it decreases cellular fitness and can cause cell death. Doxycycline is small molecule that freely moves inside the cell and associates with free rtTA forming a complex, dox-rtTA. The dox-rtTA complex then binds to the promoter of both the rtTA gene and the reporter-resistance gene, activating their gene expression and providing protection against Zeocin. However, the induced cells have a toxicity effect by squelching (Baron et al, 1997; Nevozhay et al, 2012) (see SI Section 4 for modeling toxicity effect at the cellular level).

We evolved five replicate PF yeast cell populations in seven conditions (D0Z0, D2Z2, D2Z4, D4Z2, D4Z4, D6Z2, and D6Z4). In all conditions, fitness increased for all replicates [Fig. 5(A)], similar to observation in long-term evolution experiments in bacteria (Good et al, 2017). As expected, the fitness increase (2-3 fold) was most dramatic for the non-control conditions (D2Z2, D2Z4, D4Z2, D4Z4, D6Z2, and D6Z4) compared to a smaller increase in fitness observed in the control condition (D0Z0). These results indicate that the cells adapted to the drug, and to a lesser extent, other controlled experimental conditions such as the cell culture medium, and possibly other ecological and evolutionary processes (Good et al, 2017).

**Figure 5:**
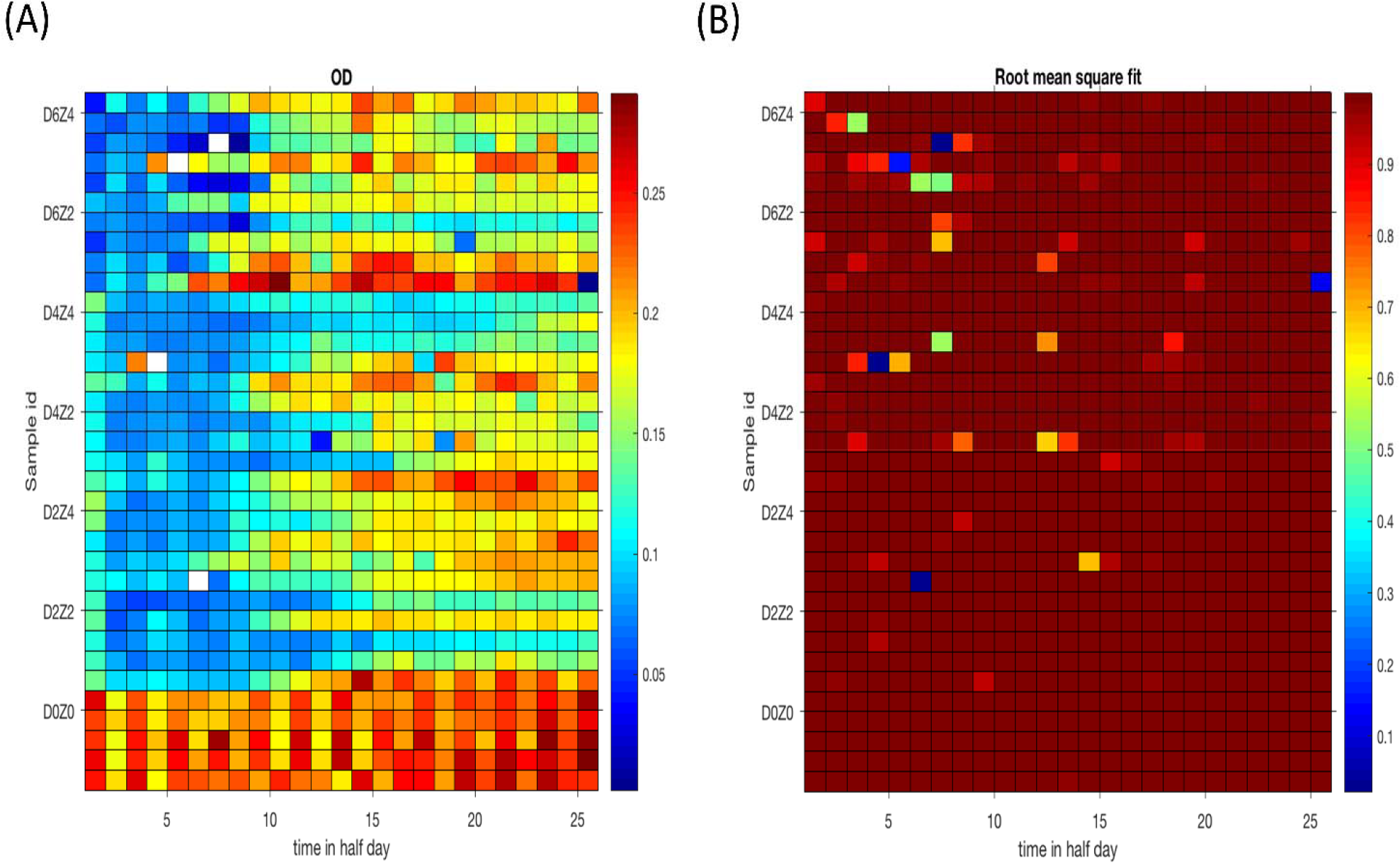
Heat map for population fitness for different combinations of inducer and drug. DiZj denotes experimental conditions, where Di and Zj indicate the concentrations of the inducer Doxycycline (μg/ml) and the drug Zeocin (mg/ml), respectively. **(A)** The color bar indicates the growth rate (fitness) of each of the colored squares, and a white square denotes a cell population crash (growth rate ≤ 0 h^−1^). **(B)** The color bar indicates the root mean square (RMS) values of a fit of an exponential function to extract population growth rate. RMS values ≪ 1 indicate a fitness swing and RMS values << 1 indicate a fitness crash due to mutation and experimental conditions.

As predicted by our model, fitness crashes occurred in experimental conditions that optimized the fitness benefit of the mutants with the time delay. Fitness crashes were observed in the D2Z4, D4Z4, and D6Z4 conditions. In agreement with model predictions [Fig. 4(C & D)], the increase in time delay at higher concentrations of Doxycycline for the Zeocin 4 mg/ml condition resulted in more fitness crashes and fitness swings in the evolving PF cell population (Fig. 5). Specifically, one fitness crash occurred in each of the D2Z4 and D4Z4 conditions, and two fitness crashes occurred in the D6Z4 condition [Fig. 5(A)]. No fitness crashes were observed in any of the Zeocin 2 mg/ml conditions, which indicates that this lower Zeocin concentration was sublethal for the mutant PF yeast cells. Similarly, an increasing number of fitness swings were observed as the Doxycycline concentration was increased from for a given Zeocin concentration [Fig. 5(B)]. As expected, no fitness crashes or fitness swings occurred in any of the D0Z0 control replicates (Fig. 5). Overall, the experimental evolution results qualitatively support our modeling predictions.

## 3 Conclusions

We show for the first time a paradoxical result that a deleterious time-delayed effect of an initially beneficial mutation can result from an interaction with gene regulatory network dynamics and cause a permanent decrease in population fitness in a constant environment. This is a novel example of how intracellular biochemical kinetics (perturbation propagating through a gene regulatory network) can affect population and evolutionary dynamics (Sanchez & Gore, 2013). This phenomenon differs from genetic drift, a process that occurs predominately in small populations which randomly alters allele frequencies and reduces genetic variability by allowing beneficial, neutral, or deleterious alleles to fix in the population, in that 1) it occurs in large populations, 2) it is repeatable and can be predicted from our mathematical models, and 3) it always negatively impacts population fitness when the time lag is sufficiently long and/or the initial fitness benefit is sufficiently large.

Using different modeling approaches, we introduced a single mutant cell into a population of wild-type cells, and traced the dynamics of the mutation by changing the time delay of mutation and the cellular fitness benefit of the mutant. This single mutation had different effects depending on the combination of the time delay and cellular fitness. For a long time delay and high mutant cell fitness, the mutant cells took over the population, as generally expected for strong selection. In this scenario, the resulting mutant population’s fitness stabilized at a lower value than the original wild-type population fitness. For a short time delay and/or low cellular fitness benefit, the fraction of mutant cells increased at an early time causing an increase in the fitness of the population, but then the cellular fitness dropped after some time and the mutant population collapsed. At intermediate levels of delay and cellular fitness benefit, the mutant cells underwent stochastic fate decision. The mutant subpopulation either swept the population, causing fitness to stabilize at a lower value than the original wild type population fitness, or it collapsed.

The evolution experiments in our study provide support for the “backwards evolution” phenomenon predicted by our models. Though all replicates increased in fitness over the course of the experiment, replicates crashed in inducer and drug where the time delays where sufficiently long and fitness benefit sufficiently large. This suggests that in these replicates the synthetic positive feedback circuit had mutated to optimize fitness [similar to what was observed previously in experimental evolution of the PF gene circuit (González et al, 2015)], which then lead to the fixation of a subsequent mutation that was ultimately deleterious and lead to the population crash. Further experimentation along with sequencing is required to ultimately confirm this hypothesis. Experimentation at a single-cell resolution using microscopy and microfluidics will also allow for a more fine-grained investigation of the interplay between gene expression and fitness dynamics.

There are other cases in nature where fitness is temporally modulated. For instance, it has been suggested that the R72 allele of the p53 tumor suppressor gene is an example of antagonistic pleiotropy, as R72 expression increases implantation of the blastocyst into the uterus while reducing the longevity in humans (Hu et al, 2008). Another example is the oscillatory predator-prey dynamics of interacting populations, such as the concentrations of *E. coli* bacteria and bacteriophage T2 (Levin et al, 1977). Furthermore, it is well known that a mutation that is beneficial in one environment may be deleterious in another environment, and that repeated population bottlenecks can drive the fixation of random deleterious mutations resulting in a decline in fitness (Beaumont et al, 2009). However, here we have demonstrated how a mutation can be initially beneficial and then become detrimental in the same environment due to the interaction of a gene regulatory network with population-level dynamics.

The findings presented in this study may have clinical relevance in conditions where mutating or inhibiting in the regulators of drug resistant states confer temporary fitness benefits followed by a fitness detriment to pathogenic cell populations. This opens new lines of investigation for treatments that could consider similar interactions of regulatory network dynamics and evolutionary dynamics to lower the fitness of pathogenic microbes or cancer cells.

## 4 Materials and Methods

### 4.1 Mathematical Models

The model for wild-type cells consists of a coupled system of ordinary differential equations (ODEs)

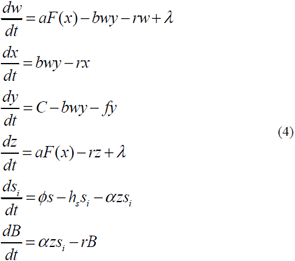

where the variables *w*, *x*, *y*, *z*, *s_i_*, and *B* are the free rtTA activator molecules, the rtTA-Doxycycline complex, the inducer Doxycycline, the free (or unbound) reporter-resistance protein molecules (GFP::ZeoR), the intracellular Zeocin antibiotic stressor, and the GFP::ZeoR-Zeocin stressor complex, respectively. The parameters *b*, *r*, *C*, and *f* are respectively the rtTA-Doxycycline binding rate, dilution rate due to cell growth of rtTA, rtTA-Doxycycline and the effector molecules, extracellular Doxycycline concentration, effective intracellular Doxycyline degradation, dilution, and outflux rate, and λ and *a* + λ are the minimum and maximum production rates of the activator and effector proteins, respectively. The parameters *s*, *ϕ*, *h_s_*, α represent the extracellular Zeocin concentration, influx rate of Zeocin across the cellular membrane, and the combined intracellular Zeocin degradation and dilution rate. See SI Section 3.1 for more details on the model and for parameter values. The response function of the protein activator is described by a Hill function

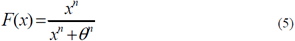

where *θ* is the dissociation constant and *n* is the Hill coefficient.

Similarly, the ODE model for mutant cells is given by

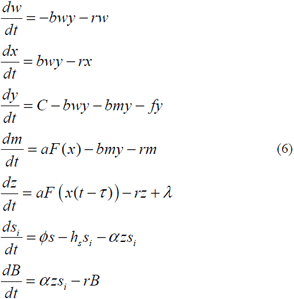

where *m* is the mutant protein and *b* is now also the binding rate between the mutant rtTA and Doxycycline. See SI for further details on the deterministic model (SI Section 3.1), as well as a full description of the stochastic models (SI Sections 3.2-3.3).

### 4.2 Experimental Methods

Well-isolated single *Saccharomyces cerevisiae* colonies carrying the genomically integrated PF gene circuit (Nevozhay et al, 2012) were picked from plates and incubated overnight in 2% glucose + SD -his -trp + ade medium at 30°C. Twelve hours later, the 1 ml cell suspensions were diluted to the concentration 1 × 10^6^ cells/ml - concentration estimated using a Nexcelom Cellometer Vision cell counter (Nexcelom Bioscience, Lawrence, MA) - in fresh SD -his -trp + ade medium supplemented with 2% galactose. Quadruplicate cultures were grown for 48 hours in the galactose medium with the appropriate concentration of Doxycycline (Acros Organics, Geel, Belgium) for each condition to allow gene expression levels to stabilize (Nevozhay et al, 2009; Nevozhay et al, 2012). Cell density was measured (Nexcelom Cellometer Vision) and resuspended to a concentration 1×10^6^ cells/ml and transferred to a 96-well microplate and placed in an Infinite M200 Pro plate reader (Tecan) at °30C. Using the Tecan plate reader, OD_600_ measurements (600 ± 9 nm, number of reads = 25) of orbitally shaken (280 rpm with an amplitude of 2 mm) 200 μl cultures were acquired every 20 minutes and cultures resuspended to an OD of 0.1 in fresh 2% galactose + SD -his -trp + ade + Doxycycline (concentration as required by experimental condition) + Zeocin (concentration as required by experimental condition) every 12 hours to keep them in log-phase growth. When a fitness swing occurred, cells were taken from the corresponding replicate in the microplate from the previous 12 hour interval in order to continue the experiment.

## Acknowledgments

This work was supported by the Laufer Center for Physical & Quantitative Biology, by the NIH Director’s New Innovator Award Program [Grant No: 1DP2 OD006481-01] and by NSF grant BIO IOS (Integrative Organismal Systems) 1021675 to G.B. D.C. was supported by an NSERC Postdoctoral Fellowship [Grant no: PDF-453977-2014]. We thank Rhys Adams and Tamás Székely for helpful discussions, and Mirna Kheir for help editing the manuscript.

## Author contributions

M.B, D.C., and G.B designed research; M.B. and D.C. performed the modeling and simulations; D.C. performed the experiments; M.B. analyzed data; and M.B., D.C., and G.B. wrote the paper.

## References

Acar M, Mettetal JT, van Oudenaarden A (2008) Stochastic switching as a survival strategy in fluctuating environments. Nat Genet 40: 471-475

Alon U (2007) Network motifs: theory and experimental approaches. Nat Rev Genet 8: 450-461

Balaban NQ, Merrin J, Chait R, and et al LK (2004) Bacterial persistence as a phenotypic switch. Science 305: 1622-1625

Balázsi G, van Oudenaarden A, Collins JJ (2011) Cellular decision making and biological noise: from microbes to mammals. Cell 144: 910-925

Baron U, Gossen M, Bujard H (1997) Tetracycline-controlled transcription in eukaryotes: novel transactivators with graded transactivation potential. Nucleic Acids Res 25: 2723-2729

Beaumont HJE, Gallie J, Kost C, Ferguson GC, Rainey PB (2009) Experimental evolution of bet hedging. Nature 462: 90-93

Bornholdt S (2001) Modeling genetic networks and their evolution: a complex dynamical systems perspective. Biol Chem 382: 1289–1299

Charlebois DA, Abdennur N, Kaern M (2011a) Gene expression noise facilitates adaptation and drug resistance independently of mutation. Phys Rev Lett 107: 218101-218101

Charlebois DA, Balazsi G, Kaern M (2014) Coherent feedforward transcriptional regulatory motifs enhance drug resistance. Phys Rev E 89: 052708

Charlebois DA, Intosalmi J, Fraser D, Kaern M (2011b) An algorithm for the stochastic simulation of gene expression and heterogeneous population dynamics. Commun Comput Phys 9: 89-112

Charlebois DA, Kaern M (2012) What all the noise is about: the physical basis of cellular individuality. Can J Phys 90: 919-923

Cohen AA, Geva-Zatorsky N, Eden E, al MF-Morgenstern e (2008) Dynamic proteomics of individual cancer cells in response to a drug. Science 322: 1511-1516

Crombach A, Hogeweg P (2008) Evolution of evolvability in gene regulatory networks. PLoS Comput Biol 4: e1000112

de Visser JAGMM, Krug J (2014) Empirical fitness landscapes and the predictability of evolution. Nature Reviews Genetics 15: 480-490

Dekel E, Alon U (2005) Optimality and evolutionary tuning of the expression level of a protein. Nature 436: 588-592

Delahodde A, Delaveau T, Jacq C (1995) Positive autoregulation of the yeast transcription factor Pdr3p, which is involved in control of drug resistance. Mol Cell Biol 15: 4043-4051

Deris JB, Kim M, Zhang Z, Okano H, Hermsen R, Groisman A, Hwa T (2013) The innate growth bistability and fitness landscapes of antibiotic-resistant bacteria. Science (New York, NY) 342: 1237435-1237435

Dykhuizen DE, Dean AM, Hartl DL (1987) Metabolic flux and fitness. Genetics 115: 25-31

Elowitz MB, Levine AJ, Siggia ED, Swain PS (2002) Stochastic gene expression in a single cell. Science (New York, NY) 297: 1183-1186

Fraser D, Kaern M (2009) A chance at survival: gene expression noise and phenotypic diversification strategies. Molecular microbiology 71: 1333-1340

Garcia-Bernardo J, Dunlop MJ (2013) Tunable stochastic pulsing in the escherichia coli multiple antibiotic resistance network from interlinked positive and negative feedback loops. PLoS Comput Biol 9: e1003229

Glanemann C, Loos A, Gorret N, Willis LB, O’Brien XM, Lessard PA, Sinskey AJ (2003) Disparity between changes in mRNA abundance and enzyme activity in Corynebacterium glutamicum: implications for DNA microarray analysis. Appl Microbiol Biotechnol 61: 61-68

González C, Ray JCJ, Manhart M, Adams RM, Nevozhay D, Morozov AV, Balázsi G (2015) Stress-response balance drives the evolution of a network module and its host genome. Mol Syst Biol 11: 827

Good BH, McDonald MJ, Barrick JE, Lenski RE, Desai MM (2017) The dynamics of molecular evolution over 60,000 generations. Nature 551: 45-50

Hasty J, McMillen D, Isaacs F, Collins JJ (2001) Computational studies of gene regulatory networks: in numero molecular biology. Nature reviews Genetics 2: 268-279

Hu W, Feng Z, Atwal GS, Levine AJ (2008) p53: A new player in reproduction. Cell Cycle 7: 848-852

Janga S, Collado-Vides J (2007) Structure and evolution of gene regulatory networks in microbial genomes. Res Microbiol 158: 787-794

Kellis M, Patterson N, Endrizzi M, Birren B, Lander ES (2003) Sequencing and comparison of yeast species to identify genes and regulatory elements. Nature 423: 241–254

Khalili S, Lin Y, Armaou A, Matsoukas T (2010) Constant number monte carlo simulation of population balances with multiple growth mechanisms. AIChE journal 56: 3137–3145

Kimura M (1962) On the probability of fixation of mutant genes in a population. Genetics 47: 713

Levin BR, Stewart FM, Chao L (1977) Resource-limited growth, competition, and predation: a model and experimental studies with bacteria and bacteriophage. Amer Natur 111: 3–24

Levy SF, Ziv N, Siegal ML (2012) Bet hedging in yeast by heterogeneous, age-correlated expression of a stress protectant. PLoS biology 10: e1001325-e1001325

Mantzaris NV (2006) Stochastic and deterministic simulations of heterogeneous cell population dynamics. Journal of theoretical biology 241: 690-706

Mantzaris NV (2007) From single-cell genetic architecture to cell population dynamics: quantitatively decomposing the effects of different population heterogeneity sources for a genetic network with positive feedback architecture. Biophysical journal 92: 4271-4288

Maynard ND, Birch EW, Sanghvi JC, Chen L, Gutschow MV, Covert MW (2010) A forward-genetic screen and dynamic analysis of lambda phage host-dependencies reveals an extensive interaction network and a new anti-viral strategy. PLoS Genet 6: e1001017

Mitrophanov AY, Groisman EA (2008) Positive feedback in cellular control systems. Bioessays 30: 542–555

Nevozhay D, Adams RM, Murphy KF, Josic K, Balázsi G (2009) Negative autoregulation linearizes the dose-response and suppresses the heterogeneity of gene expression. Proc Natl Acad Sci USA 106: 5123-5128

Nevozhay D, Adams RM, Van Itallie E, Bennett MR, Balázsi G (2012) Mapping the environmental fitness landscape of a synthetic gene circuit. PLoS Comput Biol 8: e1002480

Nevozhay D, Zal T, Balázsi G (2013) Transferring a synthetic gene circuit from yeast to mammalian cells. Nat Commun 4: 1451-1451

O’Brien EJ, Lerman JA, Chang RL, Hyduke DR, Palsson BØ (2013) Genome-scale models of metabolism and gene expression extend and refine growth phenotype prediction. Mol Syst Biol 9: 693

Peng W, Liu P, Xue Y, Acar M (2015) Evolution of gene network activity by tuning the strength of negative-feedback regulation. Nat Commun 6: 6226

Prud’homme B, Gompel N, Carroll SB (2007) Emerging principles of regulatory evolution. Proc Natl Acad Sci USA 104: 8605–8612

Rancati G, Pavelka N, Fleharty B, Noll A, Trimble R, Walton K, Perera A, Staehling-Hampton K, Seidel CW, Li R (2008) Aneuploidy underlies rapid adaptive evolution of yeast cells deprived of a conserved cytokinesis motor. Cell 135: 879-893

Sanchez A, Gore J (2013) Feedback between population and evolutionary dynamics determines the fate of social microbial populations. PLoS Biology 11: e1001547

Shen-Orr S, Milo R, Mangan S, Alon U (2002) Network Motifs in the Transcriptional Regulation Network of Escherichia coli. Nat^~^Genet 31: 64-68

van Kampen NG (1992) Stochastic Processes in Physics and Chemistry: Amsterdam: North-Holland.

Verhulst P-F (1845) Recherches mathématiques sur la loi d’accroissement de la population. Nouv Mém Acad R Sci B-lett Brux 18: 1-45

Verhulst P-F (1847) Deuxième mémoire sur la loi d’accroissement de la population Mém Acad R Sci Lett B-Arts Belg 20: 142-173

Yeh PJ, Hegreness MJ, Aiden AP, Kishony R (2009) Drug interactions and the evolution of antibiotic resistance. Nat Rev Microbiol 7: 460–466

